# Lipidated ApoE is found in nanoscale proximity to Aβ aggregates in human Alzheimer brains

**DOI:** 10.64898/2026.05.30.729004

**Authors:** Desmond Owusu Kwarteng, Rosemary J. Jackson, Tsuneo Nakajima, Kirsten Altig, Alexandra Melloni, Derek Oakley, Alberto Serrano-Pozo, Masato Maesako, Bradley T. Hyman

**Affiliations:** Alzheimer Research Unit, MassGeneral Institute for Neurodegenerative Disease, Charlestown, MA 02129, USA; Department of Neurology, Massachusetts General Hospital, Boston, MA 02114, USA; Department of Pathology, Massachusetts General Hospital, Boston, MA 02114, USA; Massachusetts Alzheimer’s Disease Research Center (MADRC), Charlestown, MA 02129, USA; Harvard Medical School, Boston, MA 02115, USA

**Keywords:** Apolipoprotein E, Amyloid beta, anionic lipids, cholesterol, FLIM-FRET, Alzheimer’s disease

## Abstract

Apolipoprotein E (*APOE*) associates with amyloid plaques (Aβ) in Alzheimer disease (AD). The ε4 allele of apolipoprotein E (*APOE*ε4) is the strongest genetic risk factor for sporadic AD and exacerbates Aβ plaque burden relative to *APOE*ε3 and *APOE*ε2. The majority of ApoE associates with multiple lipid classes to form lipoproteins both in the brain and the periphery. However, the lipidation status of Aβ plaque-associated ApoE is not yet fully defined. Here, we use fluorescence lifetime imaging microscopy coupled with Förster resonance energy transfer (FLIM-FRET) to determine the lipidation status of ApoE in plaques, as well as the nanoscale spatial proximity of ApoE and Aβ to anionic lipids and cholesterol within human AD brain tissue. We demonstrate that lipids are in close nanoscale proximity to ApoE and Aβ within Aβ plaques. Our results reveal that lipidated ApoE complexes enriched in anionic lipids and cholesterol are core constituents of AD plaques *in-situ*. We propose a pathological mechanism in which the surface presentation of anionic lipids on ApoE lipoproteins facilitates initial interaction with and subsequent aggregation of Aβ.

## Introduction

Alzheimer’s disease (AD) is characterized by deposition of amyloid beta (Aβ) which aggregates and deposits in association with multiple other proteins as senile plaques^1,2^. One of the co-depositing proteins, apolipoprotein E (ApoE), is a major genetic risk factor for sporadic AD^3,4^. *APOE*ε4 homozygous individuals are 12 to 15 times at risk of developing late-onset AD and have a high Aβ plaque deposition relative to *APOE*ε3 homozygous individuals^5,6^. In contrast, this phenotype is ameliorated in *APOE*ε2 carriers^5,6^. In animal models, *Apoe* knockout leads to a marked reduction of Aβ deposition^7,8^, whereas knock-in of human *APOE* alleles mimics the pattern of Aβ burden seen in humans, with ε4 > ε3 > ε2^9^. ApoE is known to directly interact with Aβ^10^, which is an amphipathic peptide with a hydrophobic region that likely interacts with the lipid-facing domains of ApoE^11,12^. Recent studies in mice utilizing an antibody that preferentially recognizes poorly lipidated ApoE suggest that ApoE-lipid interactions may contribute to ApoE’s impact on Aβ deposition in AD ^13,14^.

Analogous to their interactions with other anionic molecules^15,16^, ApoE isoforms exhibit distinct binding preferences with anionic phospholipids in model brain lipid membranes^17^. Notably, anionic lipids accelerate Aβ aggregation and increase their toxicity^18,19^. These results support the idea that ApoE lipoproteins enriched in diverse anionic lipids^20,21^ could provide favorable membrane electrostatic environment for Aβ attraction, binding, and aggregation. However, most studies on lipidated ApoE and Aβ aggregation have been performed *in vitro* and primarily used neutral lipids to form the lipid component of the lipoprotein, potentially limiting their relevance to the human brain context^22–24^. Hence, the lipidation status of ApoE, and the possible mediation of Aβ aggregation by anionic lipids on the ApoE lipoprotein surface in the AD brain are unknown (Figure 1).

**Figure 1:**
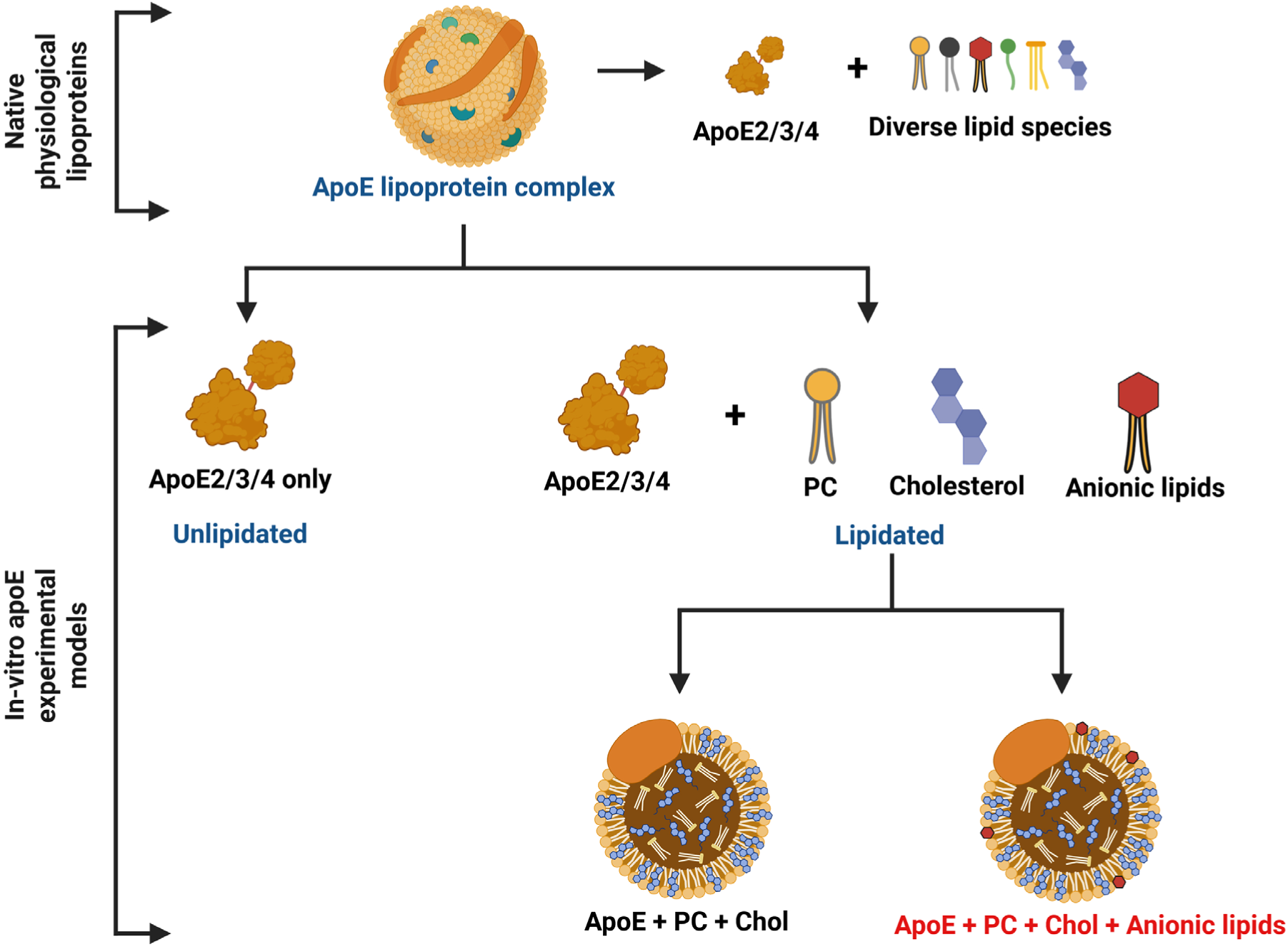
Standard *in vitro* models underrepresent the physiological rich diversity of lipid species present in native ApoE lipoproteins. Schematic of native physiological ApoE lipoprotein containing multiple lipid species compared to conventional *in vitro* models. Most lipidated ApoE investigations primarily use neutral lipids like phosphatidylcholine (PC) and cholesterol (Chol), which do make up the majority of lipid species in ApoE lipoparticles but neglect anionic phospholipids.

Herein, we tested the hypothesis that lipids associate closely with ApoE and Aβ in plaques. To test this hypothesis, we investigated the proximity of specific lipid classes (anionic phospholipids and cholesterol) to aggregated ApoE and Aβ *in situ* in postmortem human AD brains by using Florescence Lifetime Imaging Microscopy Förster-Resonance Energy Transfer (FLIM-FRET) to probe the proximity of both ApoE and Aβ to these lipids in sections from AD brain donors. We show that anionic lipids and cholesterol ester (CE) are in close proximity to both ApoE and Aβ aggregates. In addition, using a fluorescent lipid droplet dye (LD dye) that recognizes triglycerides (TG) and CE, we also confirmed a strong colocalization of neutral lipids to both ApoE and Aβ. Together, these data provide new evidence that ApoE lipoparticles enriched in both neutral and anionic lipids associate with and thus may facilitate the binding of Aβ to ApoE.

## Methods

### Human postmortem brains

Fresh frozen and formalin-fixed paraffin-embedded (FFPE) postmortem human brain tissues from the parietal region were provided by the Massachusetts Alzheimer’s Disease Research Center (MADRC) Neuropathology Core brain bank. A total of six brain donors who met the clinical^25^ and neuropathologic diagnostic criteria^26^ for Alzheimer’s disease were examined, including three *APOE*ε3 and three *APOE*ε4 homozygotes. Demographic and neuropathological characteristics, and *APOE* genotype are available in Supplementary Table 1.

### Antibodies and Reagents

#### Primary antibodies

Primary antibodies used in this study included: rabbit anti-MAP2 antibody (4542; RRID:AB_10693782; Cell Signaling Technology), rabbit polyclonal anti-IBA1 (019-19741; RRID:AB_839504; FUJIFILM Wako Pure Chemical Corporation), rabbit polyclonal anti-Apolipoprotein E (ApoE) (NBP1-31123; RRID:AB_2058116; Novus Biologicals), rabbit monoclonal anti-Aβ (clone D54D2, RRID:AB_2797642; Cell Signaling Technology), mouse monoclonal anti-Aβ (clone 6E10; 803004, RRID:AB_2715854; BioLegend), mouse monoclonal anti-phosphatidylserine (05-719, RRID:AB_309933: Millipore), mouse monoclonal anti-phosphatidylinositol (P-Z999; Echelon Biosciences), rabbit polyclonal anti-cholesterol (LS-C295824; LifeSpan BioSciences [LSBio]), and Cy3-conjugated rabbit polyclonal anti-cholesterol (LS-C701147; LSBio).

#### Validation of anti-lipid primary antibodies

Solid-phase lipid binding assays were used to determine the binding specificities of anti-cholesterol, anti-phosphatidylserine (PS), and anti-phosphatidylinositol (PI) antibodies to lipid arrays (Mega lipid strips) according to the manufacturer’s protocol. The strips contained 23 distinct lipid species immobilized at specific numbered positions (corresponding to the layout shown in Figure 1A): **(1)** Phosphatidylinositol; **(2)** Diacylglycerol; **(3)** Lysophosphatidic acid; **(4)** Phosphatidylinositol 3-phosphate; **(5)** Phosphatidic acid; **(6)** Lysobisphosphatidic acid; **(7)** Phosphatidylinositol 4-phosphate; **(8)** Phosphatidylcholine; **(9)** Lysophosphocholine; **(10)** Phosphatidylinositol 5-phosphate; **(11)** Phosphatidylethanolamine; **(12)** Sphingosine-1-phosphate; **(13)** Phosphatidylinositol 3,4-bisphosphate; **(14)** Phosphatidylglycerol; **(15)** Sphingomyelin; **(16)** Phosphatidylinositol 3,5-bisphosphate; **(17)** Phosphatidylserine; **(18)** Sulfatide; **(19)** Phosphatidylinositol 4,5-bisphosphate; **(20)** Cardiolipin; **(21)** Ceramide-1-phosphate; **(22)** Phosphatidylinositol 3,4,5-triphosphate; and **(23)** Cholesterol.

Prior to antibody incubation, the membrane strips were blocked in fatty acid-free 3% BSA in TBS-T (Tris-buffered saline containing 0.05% Tween 20) for 1 h at room temperature with gentle agitation. 8 μg/mL each of the three anti-lipid antibodies diluted in blocking buffer (fatty acid-free 3% BSA in 0.05% TBS-T) were added to different lipid strips and incubated at 4°C overnight with gentle shaking. Following overnight incubation, membrane strips were washed twice with TBS-T for 10 min each and a final 10 min wash with TBS. Strips were incubated with donkey anti-mouse IgG IRDye 800CW (1:2,000 dilution) for the anti-PI and anti-PS antibody-incubated lipid strips, and with donkey anti-rabbit IgG IRDye (1:2,000 dilution) for the anti-cholesterol antibody at room temperature for 90 mins with gentle agitation. Lipid membranes were washed twice with TBS-T for 10 min each and a final wash with TBS for 10 min. Lipid strips were imaged with Image Studio Lite system (LICOR).

#### Secondary antibodies

Fluorescently conjugated secondary antibodies used for immunostaining included: goat anti-rabbit IgG Alexa Fluor 488 (A11008; RRID:AB_143165; Thermo Fisher Scientific), donkey anti-mouse IgG Cy3 (Cat# 715-165-150, RRID:AB_2340813; Jackson ImmunoResearch Labs), donkey anti-rabbit IgG Alexa Fluor 555 (A31572; RRID:AB_162543; Thermo Fisher Scientific), donkey anti-rabbit IgG Alexa Fluor 647 (A31573; RRID:AB_2536183: Thermo Fisher Scientific), donkey anti-mouse IgG IRDye 800CW (926-32212; RRID:AB_621847; LI-COR Biosciences), and donkey anti-rabbit IgG IRDye 800CW (926-32213; RRID:AB_621848; LI-COR Biosciences).

#### Other chemicals and reagents

Additional chemicals and assay materials included: fatty acid-free Bovine Serum Albumin (BSA) (A6003; Sigma-Aldrich), Paraformaldehyde (PFA) (15710; Electron Microscopy Sciences), Phosphate-Buffered Saline (PBS) (14190144; Gibco), Tween-20 (sc-29113B; Santa Cruz Biotechnology Inc), and Mega Lipid Strips (P-6005; Echelon Biosciences). Tissue and cell preparations were mounted using Fluoromount-G Mounting Medium with DAPI (0100-20; SouthernBiotech) or a standard mounting medium without DAPI (E143470; Invitrogen).

### Immunohistochemistry and lipid histochemistry

Leica cryostat was used to slice 14-μm-thick fresh frozen sections onto a Superfrost slide. Sections were washed with TBS for 1 min and incubated in 4% PFA diluted in PBS for 15 min. Tissues were washed with TBS for 1 min and blocked in 5% fatty acid free BSA in TBS for 30 min at room temperature before overnight incubation with primary antibodies. Primary antibody dilutions were 1:250 for anti-ApoE (NBP1-31123), 1:250 for anti-Aβ (D54D2), 1:250 for anti-Aβ (6E10), 1:125 for anti-PIP (P-Z999), 1:125 for anti-PS (05-719), and 1:125 for anti-cholesterol (Cy3-conjugated, LS-C701147). Sections were washed with TBS and incubated with secondary antibodies at 1:250 dilution. Tissues for confocal imaging only (not used for FLIM-FRET) were stained with A488, A555 or A647-conjugated secondary antibodies. Sections were washed with TBS and mounted with Fluoromount-G mounting media with DAPI. For FLIM experiments, sections were incubated with A488-conjugated goat anti rabbit for ApoE and Aβ (D54D2) primary antibodies, and Cy3-conjugated donkey anti mouse antibodies for all other primary antibodies except for anti-cholesterol antibody, which was already conjugated to Cy3 secondary. Slides were then mounted with mounting media without DAPI.

To examine the potential for nonspecific staining of reagents directed to neutral lipids, we used standard FFPE sections, which were first deparaffinized in xylenes, rehydrated through a graded series of ethanol, and boiled in citrate buffer for 20 min for antigen retrieval before cooling in the citrate buffer to room temperature. These treatments would be expected to remove any neutral lipids remaining in the sections but would likely be less effective at removing polar lipids, including PS and PIP. Sections were then washed with TBS and blocked with blocking buffer for 1 h. This was followed by incubation with primary antibodies in similar steps as already outlined for the fresh frozen sections. All stains were imaged on an Olympus FV3000 laser-scanning confocal microscope.

Biotium LipidSpot^TM^ Lipid droplet Stains (LD dye) were diluted to 1x in blocking buffer. They were added to tissue sections already stained with both primary and secondary antibodies. Tissues with LD dye were incubated for 30 min at room temperature before washing twice for 10 min each with TBS. This was followed by mounting in Fluoromount-G mounting media containing DAPI.

### FLIM and FRET Analyses

#### FLIM-FRET imaging strategy

FLIM-FRET imaging and a three-component analysis that accounts for the presence of autofluorescence in human tissues were used to evaluate the spatial proximity of ApoE and Aβ proteins to lipids using similar principles as previously described^27,28^. A488-labeled ApoE and Aβ served as donor fluorophores while Cy3 served as the acceptor fluorophore to label target lipids: phosphatidylserine (PS), phosphoinositides (PIP), and cholesterol. For protein-protein interaction studies and positive controls, Cy3 was used to label the designated acceptor protein (either ApoE or Aβ).

Briefly, an Olympus FV3000 confocal microscope at 40x objective was used to identify the colocalization of A488-labeled ApoE or Aβ donors with its corresponding Cy3-labeled acceptor within Aβ plaques. Donors were excited by a chameleon Ti:Sapphire laser (Coherent Inc., Santa Clara, CA, USA) set at 850 nm and the emission was selected using the ET525/50-2p filter (Coherent Inc., Santa Clara, CA, USA). A high-speed photomultiplier tube (MCP R3809; Hamamatsu, Bridgewater, NJ) together with a time-correlated single-photon counting acquisition board (SPC-830; Becker and Hickl, Berlin, Germany) were used to record excited donor lifetimes.

#### Three-Component Lifetime Analysis

To accurately measure FRET in postmortem brain tissue sections while eliminating the interference of conspicuous autofluorescence typical of human tissues, a multi-exponential decay analysis was performed using SPC Image software v 8.9.85.0 (Becker and Hickl) as previously described. The baseline lifetimes of endogenous tissue autofluorescence (t1) were determined in unstained tissue sections for each sample. The lifetime of the A488 donor only (in the absence of Cy3 acceptor) was established. The actual donor-only lifetime (t2) was determined through a double-exponential fit where the known autofluorescence lifetime (t1) was fixed and excluded from the analysis. Ten regions of interest (ROI) lifetimes were taken from the defined plaque region following the double exponential fit. The lifetime of the donor in the presence of the Cy3 acceptor (t3) was measured by fitting the lifetimes of t1 and t2 recorded from adjacent sections of the same individual to three fluorescence decay curves. The reduced donor lifetime (t3) was recorded across ten regions of interest (ROIs), and the mean value of these measurements was used for all statistical analyses.

The FRET efficiency, E_FRET_, which is the shortening of A488 donor fluorophore lifetime in the presence of Cy3 acceptor fluorophore expressed as a percentage, was calculated as.

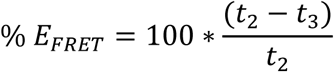

where t2 is the lifetime of the A488 donor fluorophore only (no acceptor or absence of FRET), and t3 is the A488 lifetime in the presence of Cy3 acceptor (possible FRET).

The intermolecular distances (r) between A488-labeled ApoE or Aβ and Cy3-labeled lipids were calculated using the equation:

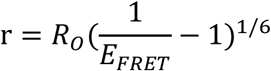

where Rₒ, the Förster radius at which 50% of energy is transferred from donor to acceptor for A488 and Cy3 donor-acceptor pairs, was assumed to be 6.75 nm^27,29^.

### Statistical Analyses

All statistical analyses were performed using GraphPad Prism (version 10; Dotmatics, Boston, MA). To assess molecular interactions, FRET parameters (donor lifetimes, FRET efficiencies, and intermolecular distances) were calculated by comparing the donor-only condition to the respective donor-acceptor conditions. Initial FRET analyses utilized ApoE-A488 as the donor against independent Cy3-labeled acceptors (cholesterol, PS, PIP, and Aβ). These measurements were conducted using AD brain tissue from three homozygous *APOE*ε3 and three homozygous *APOE*ε4 carriers, allowing for direct statistical comparison of interaction efficiencies and distances between the two genotypes. Reciprocal analyses were performed on the same tissue cohorts using Aβ-A488 as the donor against Cy3-labelled ApoE and lipid acceptors. All values are given as means and standard deviations. Data distributions were first evaluated for normality using the Shapiro-Wilk test. For continuous variables that met the assumption of normality, two-group comparisons were conducted using unpaired t-test with Welch’s correction to appropriately account for potential unequal variances. For data that violated the normality assumption, the non-parametric Mann-Whitney U test was employed. Statistical significance was defined as a two-tailed p-value < 0.05.

## Results

### Validation of poly selective anti phosphoinositide phosphate and anti-phosphatidylserine antibodies

We first determined the lipid binding specificity of the commercial poly selective anti-phosphoinositide phosphate (anti-PIP) and anti-phosphatidylserine (anti-PS) antibodies^30,31^ in a dot blot assay using strips containing twenty-three different lipid species immobilized on a membrane (Figure 2A). The positions of all lipids on the lipid strip in Figure 2A are detailed in the Methods section. From the dot-blot assay, anti-PIP antibody displayed specificity for phosphatidylinositol (spot 1 on the strip) and its seven phosphorylated derivatives (spots 4, 7, 10, 13, 16, 19, and 22). In addition, it also recognized other lipids with negatively charged phosphate or sulfate headgroups. The anti-PS antibody detected PS (spot 17) and other anionic lipids. Notably, neither antibody bound to phosphatidylcholine (spot 8), phosphatidylethanolamine (spot 11), nor to cholesterol (spot 23), which together constitute the major neutral lipid classes (∼85%) of ApoE-containing lipoproteins^20,32^ Thus, while the plasma membrane environment is highly dynamic, with variable relative abundance of lipids as well as inter and intramolecular lipid-lipid interactions impacting the physicochemical properties of lipid headgroups^33,34^, these dot-blot results indicate that the anti-PIP and anti-PS antibodies recognize distinct classes of anionic lipids and do not detect neutral lipids.

**Figure 2:**
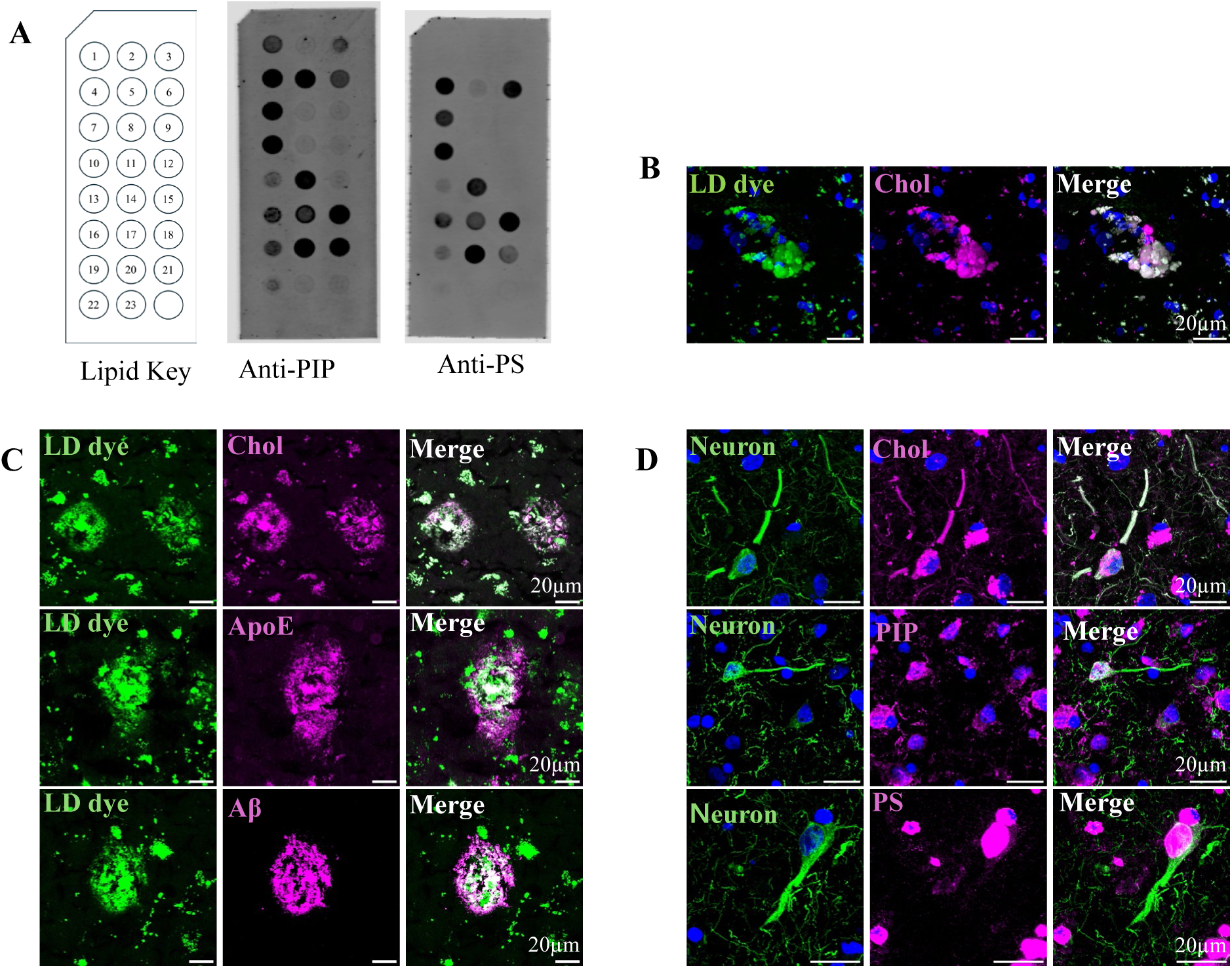
Anti-lipid antibodies demonstrate target specificity and reveal the spatial association of distinct lipid pools with neuronal and pathological markers in fresh-frozen human AD brain tissue. **(A)** Validation of anti-lipid antibody specificity using membrane lipid strips. The left panel depicts a schematic of the manufacturer-spotted lipids (distinct lipids numbered 1–23 are detailed in the Methods Section). Representative blots of identical strips (middle and right) confirm that the anti-PIP antibody specifically binds phosphatidylinositol (PI, spot 1) and its seven phosphorylated derivatives (spots 4, 7, 10, 13, 16, 19, and 22), while the anti-PS antibody binds phosphatidylserine (PS, spot 17) and related anionic lipids. **(B)** Representative confocal microscopy images of human postmortem AD brain tissue demonstrating extensive spatial colocalization (white, right) between a lipid droplet (LD) dye (LipidSpot™ 488 dye, green) and an anti-cholesterol antibody (magenta). **(C)** LD dye spatially associates with cholesterol, ApoE, and Aβ aggregates within plaque-like structures. Representative confocal microscopy images feature lipid droplets (green) visualized using LipidSpot™ 488 dye, and key pathological markers (magenta) identified via immunofluorescence (IF). In all rows, the left column displays the green lipid droplet channel, the middle column shows the magenta IF channel, and the right column displays the merged image, with white indicating spatial colocalization. *Top Row*: Co-labeling reveals that both the LD dye and anti-cholesterol antibody prominently stain dense-core, plaque-like structures. *Middle Row*: Extensive spatial overlap is observed between the neutral LD dye and anti-ApoE antibody. *Bottom Row*: Confocal images demonstrate colocalization of LD dye with Aβ aggregates. **(D)** Immunofluorescence profiling of distinct lipid targets within neurons. Representative images display co-staining of the neuronal marker MAP2 (green) alongside specific lipids (magenta): cholesterol (top row), PIP (middle row), and PS (bottom row). Merged images (far right) highlight areas of colocalization (white) between MAP2 and each respective lipid. Scale bars, 20 µm.

### Validation of anti-cholesterol antibody

Within physiological ApoE lipoprotein complexes, free cholesterol localizes to the surface monolayer, while CE are sequestered within the hydrophobic core^32^. Overall, the high abundance of these sterols (∼25 to 35%) in the lipoprotein complexes makes them suitable lipids to probe their nanoscale proximity to ApoE and Aβ aggregates^35^. We therefore assessed the lipid binding specificity of a commercial anti-cholesterol antibody using the dot-blot assay on lipid strips. The anti-cholesterol antibody failed to detect cholesterol already spotted on the lipid strip by the manufacturer and did not bind to the CE we spotted on the lipid strip (data not shown). We interpreted that the rigid nature of cholesterol and CE spotted on the membrane strip in the absence of polar lipids may form tightly packed, less fluid particles, which may not favor the binding and insertion of the anti-cholesterol antibody. Instead, we opted for immunohistochemistry, taking advantage of more fluid cholesterol packing in biological membranes due to its co-existence with other lipids, compared to 100% cholesterol spotted on the membrane strip.

A commercial Cy3-labeled anti-cholesterol antibody was used to immunostain fresh-frozen human AD postmortem brain sections counterstained with LipidSpot™ 488, a widely used dye for detecting lipid droplets because it accumulates in their TG- and CE-rich core (hereafter LD dye). Figure 2B shows extensive co-localization between the anti-cholesterol antibody and the LD dye in clusters of lipid droplets, indicating that the anti-cholesterol antibody detected CE. Notably, besides lipid droplets near the DAPI-positive nucleus of multiple cells, the LD dye also stained plaque-like objects (Figure 2C).

### Lipid droplet dye exhibits strong colocalization with both ApoE and Aβ in dense core plaques

The accumulation of lipid droplets in the brain is one of the prominent pathological features of the AD^36,37^. Recent studies have revealed that lipid droplet accumulation in glial cells of AD brains correlates with *APOE* genotype^38^. Furthermore, ApoE is found on the cytoplasmic surface of lipid droplets where it modulates the size of the droplet^38^. We used the LD dye as an indicator of neutral lipids and determined whether the dye colocalized with both ApoE and Aβ. We observed colocalizations of the LD dye to both ApoE and Aβ (Figure 2C). This finding indicates that lipidated ApoE, which physiologically contains TG and CE within its core, is associated with Aβ plaques. Alternatively, the phospholipid surfaces of lipid droplets known to contain anionic lipids may attract and aggregate ApoE and Aβ to their surfaces^39–41^. Both scenarios are consistent with several reports describing lipids associated with Aβ plaques^42–44^.

### Anionic lipids and cholesterol accumulate in neurons and microglia

The brain architecture can be viewed as a network of densely interconnected cellular membranes composed predominantly by lipids^45^. PIPs are mostly concentrated in the inner (cytoplasmic) leaflet of the plasma membranes but some reports have described them in the outer (extracellular) leaflet as well^46^. PS is abundant in the inner (cytoplasmic) leaflet but can be translocated to the outer leaflet in pathological or stressed conditions^47^. Cholesterol’s distribution on the plasma membrane is asymmetric and depends on the cell type^48,49^. Microglial and neuronal cells have cholesterol, PIPs, and PS in addition to the other lipids identified by both anionic lipid antibodies concentrated in the two leaflets of their plasma membrane^50^.

To ensure the suitability of the anti-lipid antibodies for immunohistochemistry prior to FLIM-FRET on postmortem human brain sections, we co-immunostained lipid-preserved fresh-frozen postmortem brain sections from AD donors with both anti-lipid antibodies and with neuronal-specific MAP2 antibody as well as microglial-specific anti-IBA1 antibody. Anti-PIP, anti-PS, and anti-cholesterol antibodies all exhibited widespread colocalizations with neuronal (Figure 2D) and microglial cells (Supplementary Figure 1), consistent with their relative membrane abundance. Although both anti-PIP and anti-PS antibodies detected PS in our dot-blot assay, compared to anti-PIP antibody the anti-PS antibody displayed more prominent fluorescence in AD brain sections. Thus, both immunoblotting of lipids and immunostaining of brain sections support the notion that anti-PIP and anti-PS antibodies are specific to PIPs and PS, respectively, in the complex environment of lipid membranes.

### Anionic lipids and cholesterol colocalize with ApoE in dense-core plaques

ApoE is in close proximity to and interacts with Aβ in plaques in AD brain tissues^27^. We assessed the colocalization of lipids to ApoE in lipid-preserved human AD fresh frozen sections and lipid-depleted FFPE sections. Xylene and ethanol treatment of these sections during pre-staining deparaffinization and tissue rehydration, respectively, reportedly strips lipids from the cell membrane^51,52^. We therefore used the lipid preservation differences in these two types of tissue sections as a quality control check to ensure the specificity of the lipid-binding antibodies to recognize lipids over other molecules *in situ*. Representative images of fresh frozen and FFPE sections immunostained for ApoE and lipids (PIPs, PS, and cholesterol) are shown in Figure 3 and Supplementary Figure 2.

**Figure 3:**
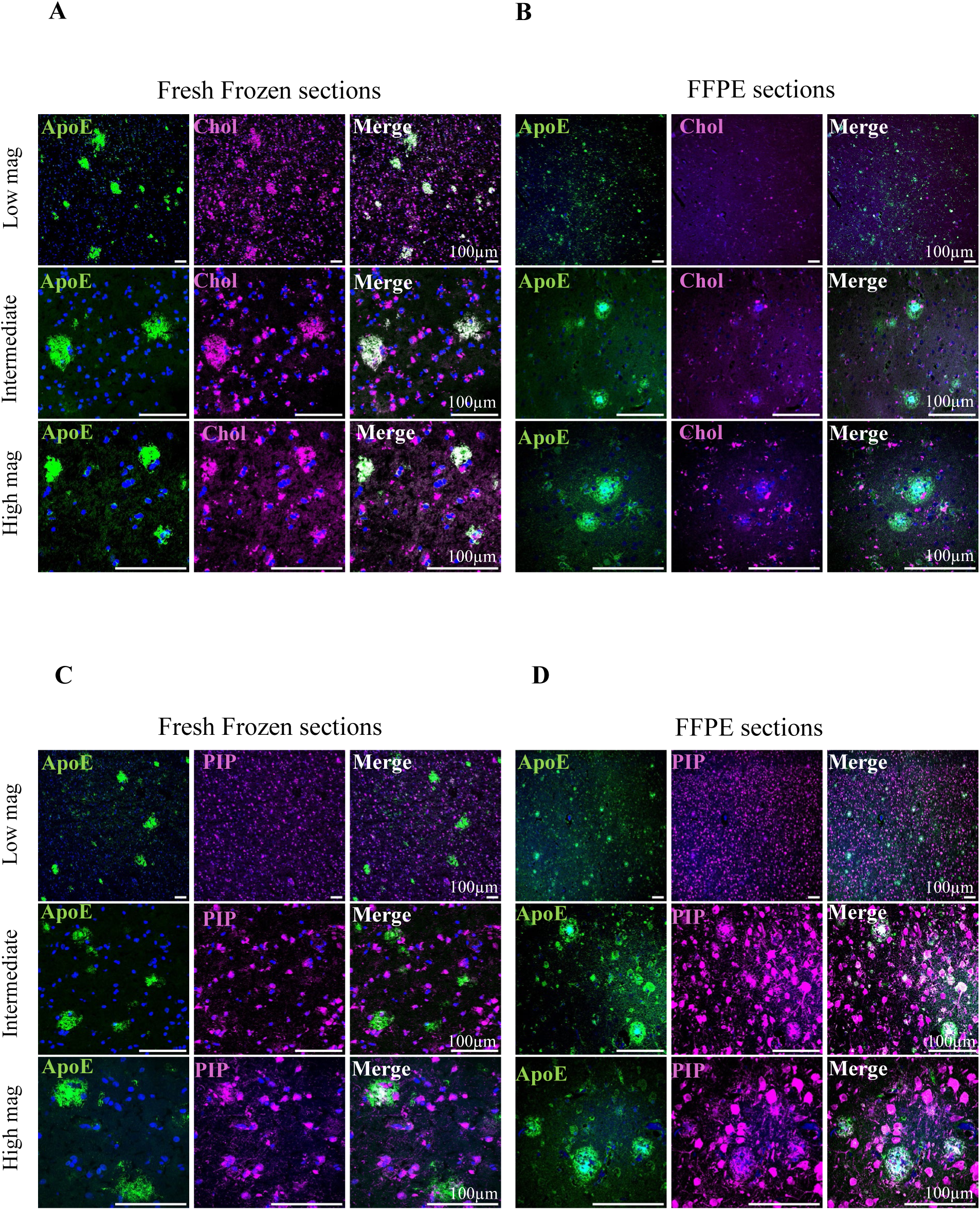
Formalin fixation and paraffin embedding severely deplete cholesterol but preserve anionic lipids in human postmortem AD brain tissue. Confocal microscopy assessing the spatial distribution and retention of ApoE and distinct lipid pools in lipid-preserved (fresh-frozen) versus lipid-depleted (FFPE) postmortem human AD brain tissue. Within each panel set, images denote ApoE (green, left), the indicated lipid target (magenta, middle), and the merged channel highlighting colocalization (white, right). Rows represent increasing magnifications from top (low power) to bottom (high power). **(A)** Co-staining for ApoE and cholesterol in fresh-frozen sections, demonstrating robust retention of the cholesterol signal. **(B)** Co-staining for ApoE and cholesterol in FFPE sections, showing severe depletion of the cholesterol signal compared to fresh-frozen tissue. **(C)** Co-staining for ApoE and phosphatidylinositol phosphates (PIP) in fresh-frozen sections. **(D)** Co-staining for ApoE and PIP in FFPE sections. Unlike cholesterol, PIP immunoreactivity remains highly preserved following formalin fixation and paraffin embedding. Scale bars, 100 µm.

Remarkably, all anti-lipid antibodies colocalized with ApoE in fresh frozen sections (Figures 3A, 3C, Supplementary 2C). Both PIPs and PS showed colocalization with aggregated ApoE at spatially restricted sites. By comparison, cholesterol showed more extensive colocalization with

ApoE in fresh frozen sections, consistent with its relatively higher presence both in the inner core and outer layer of ApoE-containing lipoproteins (Figure 3A). In the FFPE sections, as expected, there were pronounced reductions in cholesterol colocalization with ApoE relative to fresh frozen sections (Figures 3A and 3B). However, anti-PIP and anti-PS antibodies exhibited comparable intensities in areas of overlap with ApoE within dense core Aβ plaques in both tissue types. They additionally had enhanced fluorescence signals around cell bodies or processes (Figure 3D, and Sup. Figure 2D) compared to fresh frozen tissues (Figure 3C, and Sup. Figure 2C). These observations are likely attributable to the depletion of cholesterol (and neutral lipids) from the membranes during xylene and ethanol treatment in the deparaffinization and tissue rehydration procedures and / or to the antigen retrieval step of immunohistochemistry with FFPE sections. These data agree with earlier reports of the removal of neutral lipids over tightly bound protein and phospholipids interactions in FFPE sections^51,52^.

Taken together, our validation of anti-lipid antibodies via immunoblotting and immunohistochemistry demonstrates their specificity for specific lipid classes and supports their application for FLIM-FRET experiments.

### FRET positive controls validate maximal energy transfer and close proximity of donor-acceptor pairs

Using already established A488-Cy3 donor acceptor pairs, we determined the suitability of C-terminus-specific anti-ApoE and anti-Aβ antibodies for the FRET investigations. As a positive control for the ApoE FRET, we immunolabeled ApoE with both donor and acceptor fluorophores. Confocal and corresponding FLIM images are shown in Figure 4. There was significant shortening of ApoE-A488 donor lifetimes in the presence of Cy3 acceptors compared to negative controls (ApoE-A488 donor only) illustrated in Figure 5A. Similar experiments with the anti-Aβ antibody clone D54D2 yielded comparable significant lifetimes for positive and negative controls (Supplementary Figures 3 and 4).

**Figure 4:**
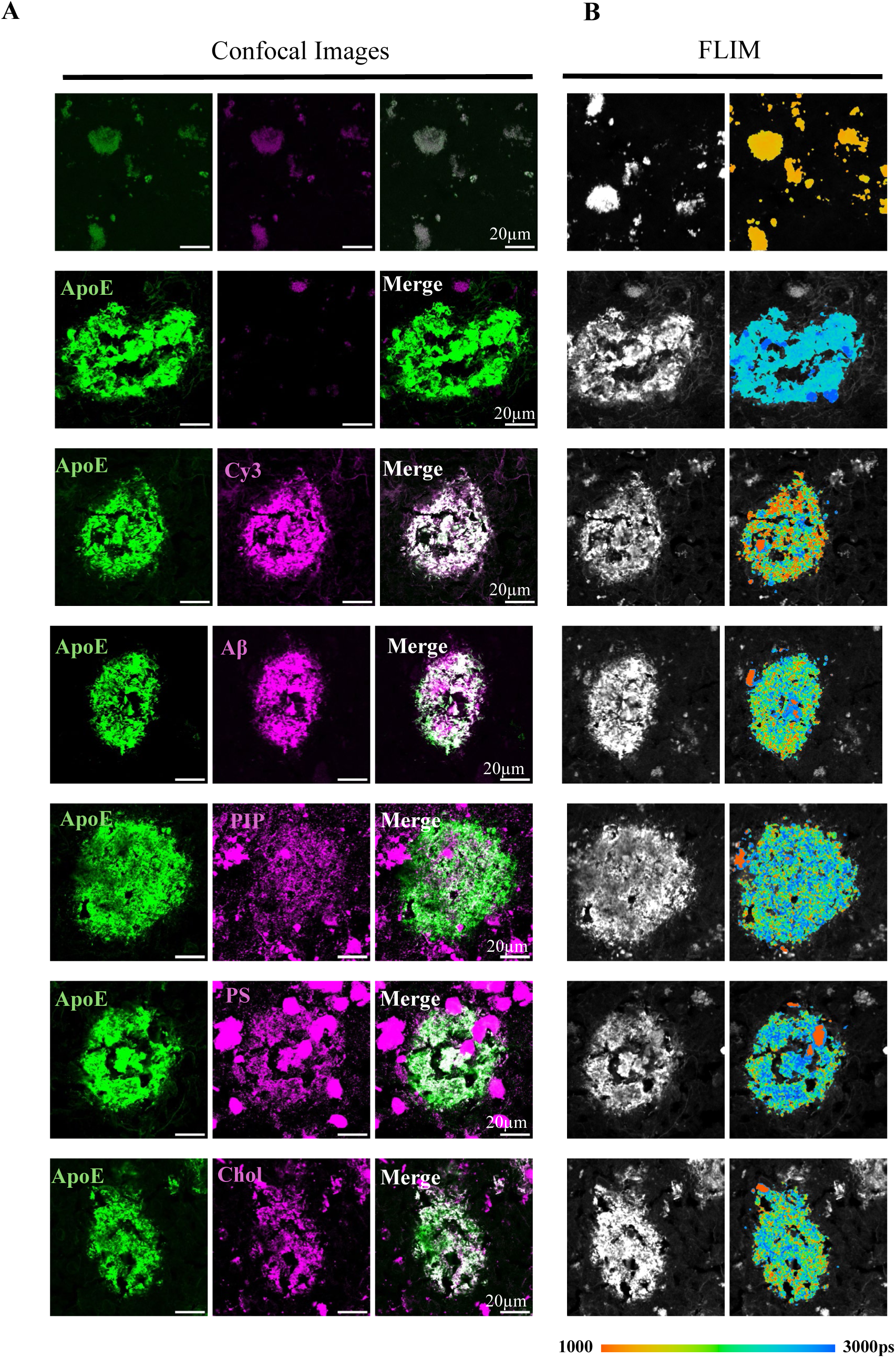
FLIM-FRET analysis reveals significant quenching of ApoE donor lifetimes by anionic lipids, cholesterol, and Aβ in human AD brain tissue. Standard image layout for all experimental conditions. **(A).** The first three columns display conventional confocal imaging of the ApoE donor (green), the indicated Cy3 acceptor (magenta), and the merged channels (white) **(B).** Subsequent columns display the corresponding FLIM intensity and pseudo-colored FLIM lifetime images. Experimental conditions and established FRET pairs. *Top Row:* Unstained human AD brain tissue to measure intrinsic autofluorescence and establish the baseline reference lifetime. *Second Row:* ApoE donor-only condition establishing the unquenched baseline fluorescence lifetime. *Third Row:* Positive FRET control utilizing ApoE labeled with a secondary antibody dual-tagged with both Alexa Fluor 488 and Cy3 to demonstrate maximal lifetime quenching. *Rows 4–7:* Experimental FRET conditions demonstrating close nanometer-scale interactions between the ApoE donor and distinct Cy3-labeled acceptors: Aβ (fourth row), PIP (fifth row), PS (sixth row), and cholesterol (bottom row). Scale bars, 20 µm. Quantified fluorescence lifetimes (picoseconds) derived from these images (brain donor ID 2219) are summarized in Table 1. This table also summarizes the ApoE FRET lifetimes across the remaining samples.

**Figure 5:**
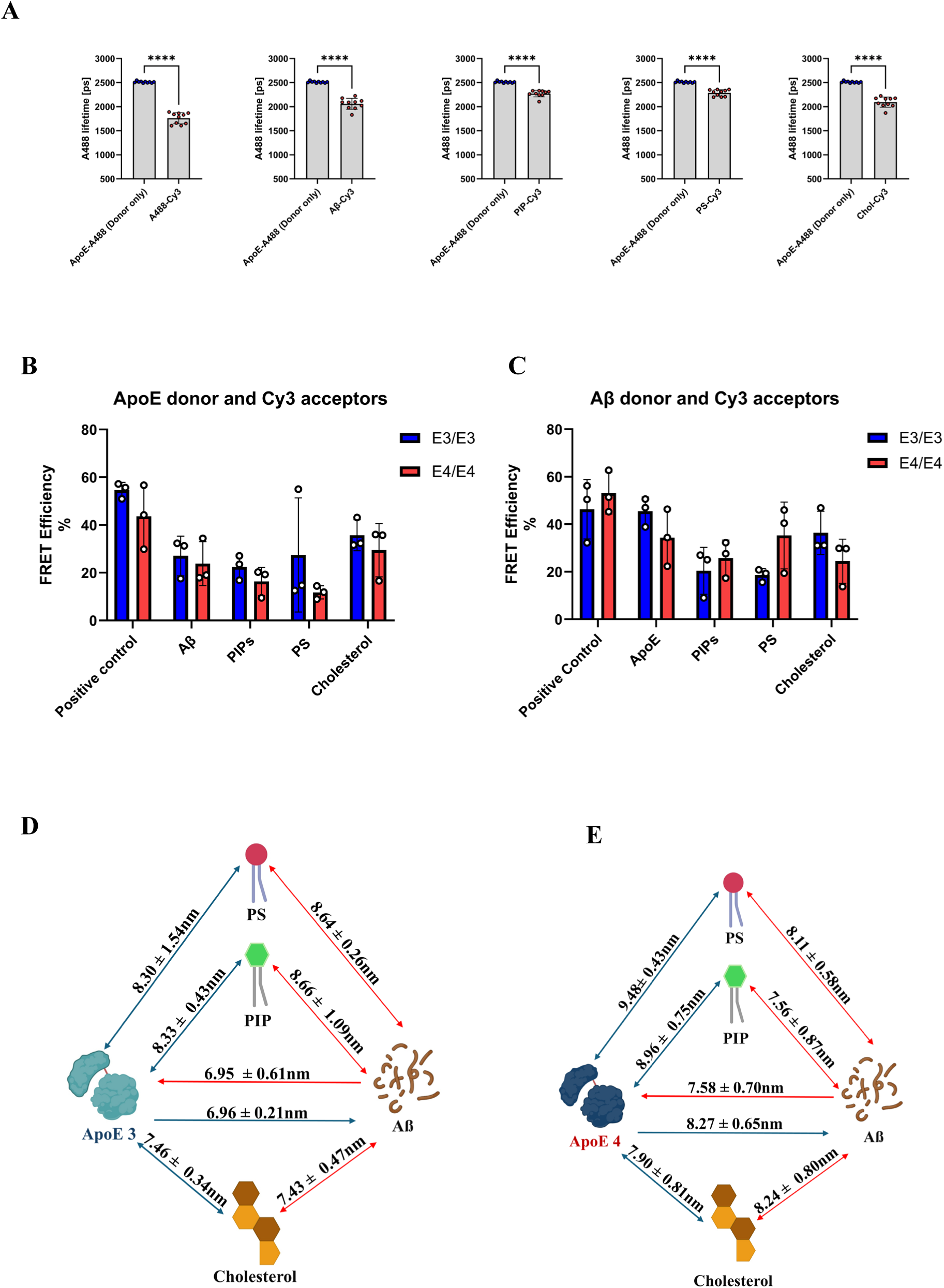
FLIM-FRET quantification reveals close interactions of ApoE-Aβ and the individual associations of ApoE and Aβ with lipid pools in *APOE*ε3 and *APOE*ε4 homozygotes. **(A)** Quantification of ApoE donor fluorescence lifetimes. Bar graphs compare the unquenched baseline lifetime of the ApoE donor against five specific Cy3-labeled acceptor conditions: (1) dual-labeled positive control, (2) Aβ, (3) PIP, (4) PS, and (5) cholesterol. Significant reductions in donor lifetime indicate closer nanoscale intermolecular proximity. **(B)** Calculated FRET efficiencies for the ApoE donor paired with specific Cy3-labeled acceptors. Data are stratified by patient’s *APOE* genotype, comparing the *APOE*ε3 (blue) and *APOE*ε4 (red) homozygotes. **(C)** Calculated FRET efficiencies for the Aβ donor paired with specific Cy3-labeled acceptors. Data are similarly stratified by *APOE* genotype as described in (B). Efficiencies in (B) and (C) were derived from the measured donor lifetimes presented in (A) and Supplementary Figure 4. **(D)** Estimated intermolecular FRET distances (nm) defining the spatial proximity between ApoE and Aβ donors and their respective acceptors (ApoE or Aβ, PIP, PS, and cholesterol) in *APOE*ε3 homozygotes. **(E)** Estimated intermolecular FRET distances (nm) defining the spatial proximity between ApoE and Aβ donors and their respective acceptors (ApoE/Aβ, PIP, PS, and cholesterol) in *APOE*ε4 homozygotes. Distances in (D) and (E) were derived from the respective FRET efficiencies calculated in (B) and (C). In figures (D) and (E), ApoE donors are represented by blue lines connecting to the acceptor molecule, and Aβ donors are shown by red lines. For all panels, error bars represent standard deviation (s.d.). Statistical significance was determined using t-test with Welch’s correction for uneven variances (****p < 0.0001).

**Table 1:** Summary of ApoE-A488 donor lifetimes (picoseconds) across various FRET-acceptor pairs. **Fluorescence lifetimes of ApoE-A488 donors.** Lifetimes (ps) are reported for donor-only controls, positive controls, and experimental groups paired with Cy3-labeled acceptors. Data are presented as mean lifetime values ± SD derived from independent experimental samples (N = 3). Statistical significance (p < 0.0001 for all samples) was determined by comparing donor lifetimes in FRET pairs against donor-only controls, with analysis performed on 10 Regions of Interest (ROIs) per sample.

| | *APOE*ε4 homozygotes | | | *APOE*ε3 homozygotes | | |
| --- | --- | --- | --- | --- | --- | --- |
|  | 2219 | 1707 | 1967 | 2302 | 2594 | 2207 |
| **Cy3 acceptors** |  |  |  |  |  |  |
| ApoE (donor only) | 2512 ± 18 | 2557 ± 22 | 2452 ± 16 | 2387 ± 28 | 2344 ± 50 | 2365 ± 75 |
| Positive Control | 1761 ± 116 | 1426 ± 120 | 1061 ± 43 | 1021 ± 23 | 1149 ± 54 | 1047 ± 42 |
| Aβ | 2060 ± 115 | 1677 ± 145 | 1986 ± 90 | 1614 ± 262 | 1933 ± 88 | 1625 ± 112 |
| PIP | 2273 ± 68 | 2064 ± 148 | 1954 ± 116 | 1740 ± 301 | 1792 ± 106 | 1966 ± 86 |
| PS | 2287 ± 67 | 2253 ± 96 | 2097 ± 66 | 1073 ± 232 | 2050 ± 92 | 2015 ± 132 |
| Cholesterol | 2095 ± 107 | 1642 ± 82 | 1568 ± 78 | 1365 ± 275 | 1586 ± 88 | 1617 ± 165 |

We then estimated FRET efficiency, which is the proportion of interacting donor-acceptor pairs and is known to be influenced by heterogenous labeling of aggregated proteins and uneven molecular orientations of donor-acceptor pairs. The average FRET efficiencies for the ApoE positive control were 55% and 43% in *APOE*ε3 and *APOE*ε4 homozygotes, respectively (Figures 5B and 5C), while those for the Aβ positive control were 46% and 53% in *APOE*ε3 and *APOE*ε4 homozygotes, respectively (Figures 5B and 5C).

We also calculated the FRET distance, which gives an estimate of the intermolecular proximity between donor and acceptor for the positive controls. The combined average estimated FRET distances for both ApoE and Aβ positive controls were 6.75 nm in *APOE*ε3 homozygotes and 6.85 nm in *APOE*ε4 homozygotes. There were no significant differences in the FRET efficiencies and distances for both ApoE and Aβ between the two *APOE* genotypes. These experiments validated the suitability of both anti-ApoE and anti-Aβ antibodies with A488 donor and Cy3 acceptor fluorophore pairs to be used in subsequent FRET experiments. Furthermore, they set the maximal baseline for expected FRET efficiencies and distances for A488-Cy3 donor-acceptor pairs in these experimental conditions.

### Significant reduction of ApoE and Aβ donor lifetimes in reciprocal FRET donor-acceptor pairings

We previously demonstrated that ApoE is closely associated with Aβ in human AD tissues^53^, and that the proportion of interacting pairs as well as the proximity of Aβ to ApoE are influenced by both ApoE’s N- or C-terminus and its isoform^27^. Due to the differences in properties of antibodies used in previous and current studies (monoclonal primary antibodies directly tagged with the Cy3 fluorophore vs. untagged monoclonal/polyclonal primary antibodies), and in order to standardize our experimental conditions, we first ascertained the proximity of ApoE as donor to Aβ and vice versa. Most importantly, the estimated FRET efficiencies and distances for ApoE and Aβ pairs will serve as relevant physiological controls for lipid FRET experiments in this study, as lipids within the ApoE lipoprotein environment may be most likely to interact with Aβ.

The FRET lifetimes, efficiencies, and distances of ApoE as donor to Aβ as acceptor and vice versa were determined for both ApoE isoforms. Confocal and FLIM images for ApoE-A488 and Aβ-Cy3 are shown in Figure 4. The corresponding lifetime, FRET efficiency, and FRET distance graphs are shown in Figures 4 and 5. The corresponding images and graphs for Aβ-A488 and ApoE-Cy3 are found in Supplementary Figures 3 and 4.

There were significant reductions in the lifetimes of both ApoE-A488 and Aβ-A488 as donors regardless of the *APOE* genotype. Although there were no significant differences in both ApoE and Aβ interacting pairs as estimated with FRET efficiency, a relatively higher proportion of Aβ interacted with ApoE3 than with ApoE4 independently of ApoE and Aβ donor-acceptor pairing orientations. There was no statistically significant difference in the estimated FRET distances from ApoE3’s C-terminus to Aβ (6.96 nm) compared to ApoE4’s C-terminus to Aβ (8.27nm) (Figures 5D and 5E). These findings were maintained independently of A488-Cy3 donor-acceptor orientations.

### Anionic lipids are closely associated with ApoE and Aβ in plaques as detected by FRET interactions *in situ*

ApoE isoforms loaded with varying concentrations of anionic lipids may provide a surface for attraction, binding, seeding, and growth of Aβ oligomers and fibrils. In this scenario, anionic lipids would be found in nanoscale proximity to both ApoE and Aβ. Confocal and corresponding FLIM images demonstrating the colocalization of ApoE with anionic lipids (PS and PIP) are shown in Figure 4. The identical imaging panel for Aβ is provided in Supplementary Figure 3. In both *APOE*ε3 and *APOE*ε4 homozygotes, the lifetime of ApoE-A488 was significantly reduced in the presence of Cy3-labeled anti-anionic lipid antibodies (Figure 5A), with similar reductions observed for Aβ (Supplementary Figure 4). The estimated FRET efficiencies and distances are detailed in Figures 5B–C and 5D–E, respectively.

In both *APOE*ε3 and *APOE*ε4 homozygotes, the average percentage of anionic lipids interacting with ApoE (20%) was highly comparable to the fraction of ApoE interacting with Aβ (25%) (Figure 5B). A similar interaction rate was observed between anionic lipids and Aβ (24%). However, the reciprocal measurement of Aβ interacting with ApoE was notably higher at 40% (Figure 5C). This variance may be attributed to the use of an alternative Aβ antibody, which was required for this specific FRET pair to prevent species cross-reactivity with the rabbit-derived ApoE antibody. Overall, because FRET requires strict nanometer-scale proximity, these findings indicate that anionic lipids clustered around the C-terminus of ApoE in the lipoprotein particles interacted with Aβ.

The average distance from ApoE3 to anionic lipids was estimated to be 8.32 nm and that of ApoE4 to anionic lipids was 9.22 nm (Figures 5D and 5E). Based on electron microscopy measurements of immunopurified ApoE lipoparticles, the diameters of ApoE from human CSF samples^54^, as well as ApoE3 and ApoE4 secreted by primary mouse astrocytes^55^, range from 8 to 20 nm. The estimated FRET distances between ApoE and anionic lipids are therefore well within the expected circular chord of a lipoprotein particle, which suggests that the detected FRETing lipids are constituents of the ApoE lipoprotein. Ultimately, the estimated FRET parameters indicate that anionic lipids are in nanoscale proximity to both ApoE and aggregated Aβ.

### Cholesterol/cholesterol esters are closely associated with ApoE and Aβ in plaques as detected by FRET interactions *in situ*

CE has distinct biophysical properties (charge and structure) compared to anionic lipids^33,34,56^. It is a prominent lipid component that exists in higher proportions in physiological ApoE lipoprotein complex and has been heavily used in experimental lipidated ApoE models^16,22,24^. We took advantage of its unique features and used CE as alternate lipid probe (acceptor) to determine the proximity of neutral lipids to ApoE and Aβ in AD brains. Confocal and FLIM images for ApoE are shown in Figure 4 and those for Aβ are in Supplementary Figure 3. The estimated FRET distance between ApoE3 and CE was 7.46 nm and its FRET efficiency was 36%, while the distance between ApoE4 and CE was 7.90 nm with an estimated FRET efficiency of 30% (Figure 5). Comparable FRET efficiencies and distances were obtained for CE and Aβ for both ApoE isoforms (Figure 5). There were no significant differences in either FRET parameter for the interactions of ApoE and Aβ with CE between the ApoE isoforms.

There was a slightly higher proportion of interacting CE molecules with both ApoE and Aβ and their proximities to each other were apparently shorter compared to anionic lipids for both ApoE isoforms. However, this calculation must be taken in the context of different experimental conditions: we used a directly Cy3-labeled anti-cholesterol antibody for the FLIM-FRET experiments while both primary anti-anionic lipid antibodies were immunolabeled with secondary antibodies conjugated with Cy3 fluorophore, leading to different assumptions when calculating average distances. Nonetheless, the data show that CE is also closely associated with ApoE and Aβ within plaques.

## Discussion

In this study, we have used FLIM FRET to demonstrate that lipids, both cholesterol and anionic lipids, are found in close proximity to both ApoE and Aβ in human AD brains. FLIM is advantageous over fluorescence intensity-based FRET measures because the fluorescent lifetime is inherently independent of the amount of fluorophore present, and because autofluorescence (which is prominent in aged human brains) gives rise to a characteristic short lifetime signal that can be mathematically accounted for^57^. The estimated FRET distances were similar for both anionic lipids and cholesterol to ApoE as well as to Aβ in *APOE*ε3 and *APOE*ε4 homozygotes. The estimated FRET efficiencies showed that similar proportions of ApoE molecules that are lipid associated also interacts with Aβ *in situ*. Furthermore, we have also established colocalization of both ApoE and Aβ with a lipid droplet dye that recognizes lipid droplets containing TG and CE. Together, our findings suggest that ApoE lipoprotein complexes containing physiologically diverse lipids including neutral and anionic lipids are found in nanoscale proximity to and interact with Aβ peptides within plaques.

ApoE forms complexes with Aβ in mouse and human AD brains^58,59^. Research into the isoform specific effects of ApoE on Aβ aggregation using experimental reconstituted lipidated and unlipidated ApoE have yielded diverging results and conclusions^22,23,60–62^. Unlipidated ApoE and artificially lipidated ApoE lipoprotein formed by using ‘standard model lipids’, PC and cholesterol fail to produce the isoform-specific and stable complexes of ApoE-Aβ aggregates^22,60,63^. This is in sharp contrast to research demonstrating that both ApoE isolated from postmortem human brains and ApoE lipoproteins secreted in cell culture media form stable, aggregated complexes with Aβ, faithfully recapitulating the aggregates found in human AD brains^4,63,64^. What accounts for the differences in the functional effects of ApoE isoforms and of unlipidated, artificially lipidated, and physiologically lipidated ApoE lipoproteins?

ApoE is secreted by astrocytes, microglia or stressed neurons^65^ in an inactive poorly lipidated form and is lipidated through an ABC Cassette Transporter A1 (ABCA1) in a phosphatidylinositide-requiring process^46^. ABCA1-deficient mice have ∼80% reduction in the levels of ApoE in both CSF and brain^66^ suggesting that proper ApoE lipidation is required to maintain its stability and, subsequently, its physiological functions. In addition to the requirement of phosphatidylinositide in ApoE lipidation, ApoE isoforms have been shown to have differential interactions with both phosphatidylinositol 4,5-bisphosphate (PIP2) and phosphatidylinositol 3,4,5-trisphosphate (PIP3) in brain lipid environments^17^. This suggests that ApoE proteins have a high probability of being attracted to different anionic lipid-rich portions in the membrane in an isoform-dependent manner (E4 > E3 > E2)^67^. ApoE variants would not only be enriched in PC and cholesterol as mostly depicted in experimental models. Instead, they would have varied anionic lipid loads based on their overall charge in addition to a broad lipid repertoire (Figure 1)^20^. Diverse lipid species including phosphatidylinositides and other anionic lipids are reported in lipidomic analyses of ApoE lipoprotein immunoprecipitated from cultured cells, mouse models, and human postmortem AD samples^20,67,68^. Indeed, multiple studies report disruption of PIP2 levels in the AD brain^69–71^. Reflecting this close biological relationship, our findings demonstrate that both anionic lipids and cholesterol are present in nanoscale proximity to aggregated ApoE, although in the vicinity of a plaque, in this study with modest sample size we do not observe differential interactions based on ApoE isoform. Colocalization of lipid droplet dye to ApoE also denotes that the core of aggregated ApoE in Aβ plaques contain the neutral lipids TG and CE.

The Aβ peptide possesses an overall net negative charge^72^. However, molecular dynamics simulations reveal that its interaction with membranes is mediated by initial electrostatic contacts between the peptide’s cationic lysine residue(s) and anionic phosphate headgroups, followed by subsequent hydrophobic interactions that result in strong peptide-lipid binding^73–75^. Earlier work by Chauhan et al^19^ showed that Aβ aggregation is immensely increased in the presence of anionic phospholipids over neutral, zwitterionic, and anionic non-phospholipids. Furthermore, lipids and Aβ compete for the same binding spot in ApoE^76^. Collectively, these seminal studies indicate that Aβ would favorably interact with anionic lipids contained in the lipidated ApoE lipoprotein particle over ApoE protein itself. They corroborate previous studies demonstrating that Aβ forms stable complexes with ApoE and preferentially co-migrates with larger, lipid-rich, HDL-like ApoE particles^77,78^. Additionally, these findings are consistent with reports of lipids found within Aβ plaques^43,44,79^ as well as evidence demonstrating that Aβ oligomers interact with plasma membranes to induce pores in them^80^.

Lipid free (unlipidated) experimental ApoE models self-aggregate *in vitro* in an isoform dependent manner^81^, however, self-aggregation and charge variations of unlipidated ApoE2/3/4 do not directly exert any variant specific influence in their binding to and aggregation of Aβ as has been shown in multiple studies^22,23,60^. These experimental models result in the production of a short-term ApoE-Aβ complex. Tokuda et al^82^ demonstrated that delipidating cell secreted ApoE not only decreases its binding affinity for Aβ but also eliminates the isoform-specific nature of their interaction.

In contrast to this lipid-free state, ApoE proteins lipidated with PC and cholesterol are shielded from self-aggregation^81^. In alignment with Aβ’s aggregation effect by PC and cholesterol, experimental PC and cholesterol lipidated ApoE isoform models only minimally accelerate aggregation of Aβ^22,60^. PC and cholesterol are structurally supporting and neutral lipids that are devoid of the physicochemical properties needed for electrostatic attraction and stable complex formation with most proteins, including Aβ^83,84^ (Figure 6). ApoE lipoprotein complexes containing negatively charged phospholipids and sulfatides provide an electrostatic surface that enhance complex formation of ApoE with Aβ and its subsequent aggregation^67,75,85,86^. Based on ApoE2/3/4 charge-dependent interactions with anionic molecules, we propose similar charge-mediated effects on binding of nascent ApoE isoforms to anionic lipids during their lipidation. These subsequently modulate the isoform specificity of ApoE-Aβ aggregation illustrated in Figure 6. The current data thus emphasize the potential role of anionic lipids as key components of lipidated ApoE-Aβ interaction in senile plaques, raising the possibility that anionic lipids facilitate ApoE-Aβ complex formation.

**Figure 6:**
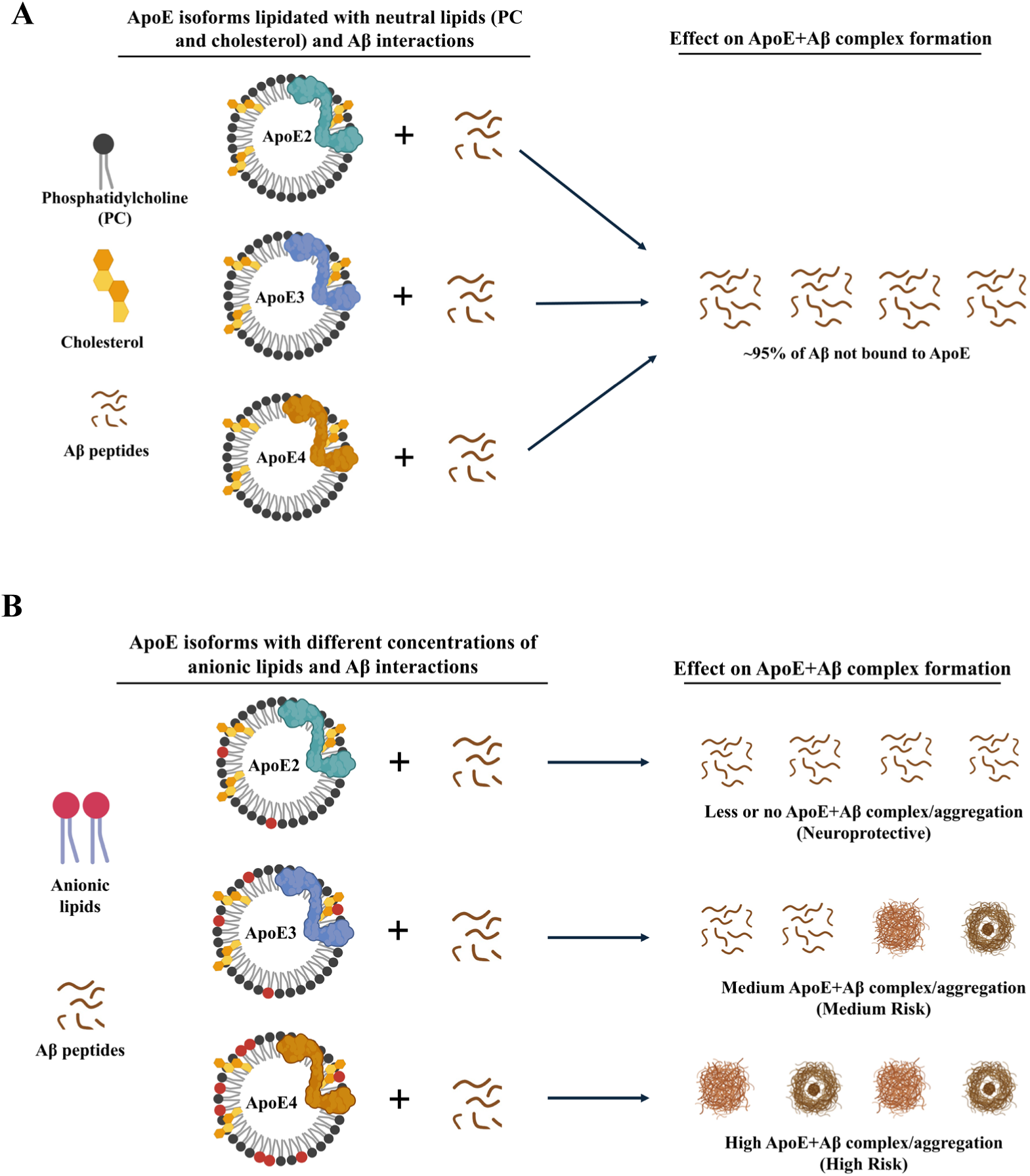
Current experimental conditions used in ApoE + lipid mediated Aβ interactions vs. proposed charge-mediated ApoE isoform-dependent differential recruitment of anionic lipids leading to differences in Aβ interaction, aggregation and AD risk. Schematic representation summarizing the differential effects of neutral versus anionic lipids on the formation of ApoE-Aβ complexes **(A) Neutral Lipid Environments:** A conceptual model depicting the three major ApoE isoforms (ApoE2, ApoE3, and ApoE4; top to bottom) lipidated with standard neutral liposomes composed of phosphatidylcholine (PC) and cholesterol. In the absence of strongly charged lipid anchors, the subsequent interaction between these lipidated ApoE lipoprotein complexes and Aβ peptides is highly inefficient, resulting in restricted complex formation (< 5% binding) as reported in most biophysical investigations. **(B) Anionic Lipid-Mediated Complex Formation:** Proposed mechanism of charge-dependent ApoE-Aβ interaction driven by negatively charged lipids. The schematic illustrates how strongly anionic lipids are differentially captured by the distinct positive charge profiles of ApoE2, ApoE3, and ApoE4. This differential capture of anionic lipids fundamentally alters the surface charge of the ApoE-lipid complex, thereby mediating and stabilizing the tight interaction with Aβ. This proposed charge-dependent mechanism visually corroborates the tight nanoscale interactions observed between ApoE, Aβ, and targeted anionic lipids in the human brain tissue FLIM-FRET data.

## Data availability

Deidentified data will be made available upon request.

## Acknowledgements

We are grateful to patients and families involved in research at the Massachusetts Alzheimer’s Disease Research Center.

## Funding

The authors gratefully acknowledge the financial support of the FTF Foundation (BTH), the Massachusetts Alzheimer Disease Research Center (NIH/NIA P30AG062421, BTH and AS-P), the Dreyfoos Fund for Alzheimer Research (BTH and AS-P), and the Alzheimer’s Association APOE Biology in Alzheimer’s Grant (ABA-25-1373553, AS-P).

## Competing interests

Dr Hyman owns stock in Novartis; he serves on the SAB of Dewpoint and has an option for stock. He serves on a scientific advisory board or is a consultant for AbbVie, Arbor Bio, Argo, Arvinas, BioClec, Biogen, BMS, Cell Signaling, Cure Alz Fund, CurieBio, Dewpoint, Eisai, Etiome, Pfizer, Sanofi, Takeda, TD Cowen, Vigil, Violet, Voyager, WaveBreak. His laboratory is supported by research grants from the National Institutes of Health, Cure Alzheimer’s Fund, Tau Consortium, and the JPB Foundation – and sponsored research agreement from Abbvie and Sanofi. He has a collaborative project with Biogen and Neurimmune. These interests were reviewed and are managed by Massachusetts General Hospital and Partners HealthCare in accordance with their conflict-of-interest policies.

AS-P has a material transfer agreement with Ionis Pharmaceuticals, Inc. and received research funds from AbbVie, Inc.

## Supplementary material

Supplementary material is available online.

