## Supplementary data for "Lipidated ApoE is found in nanoscale proximity to Aβ aggregates in human Alzheimer brains"

**Supplementary Table 1: Subject demographics and neuropathological profiles of the fresh-frozen tissue cohort. Data includes genotype, age at death, sex, post-mortem interval (PMI), and Braak stage to contextualize the FLIM-FRET microscopy findings in human AD and control samples.**

| Case Number | *APOE* Genotype | Age at Death | Sex | Braak NFT Stage | PMI (h) |
|---|---|---|---|---|---|
| 2302 | ε3/ε3 | 66 | Female | VI | 14 |
| 2207 | ε3/ε3 | 83 | Male | VI | 5 |
| 2594 | ε3/ε3 | 69 | Male | VI | 48 |
| 1797 | ε4/ε4 | 82 | Male | V | 18 |
| 1967 | ε4/ε4 | 70 | Male | V | 36 |
| 2219 | ε4/ε4 | 75 | Male | V | 10 |

Abbreviations: NFT = neurofibrillary tangle. PMI = postmortem interval.

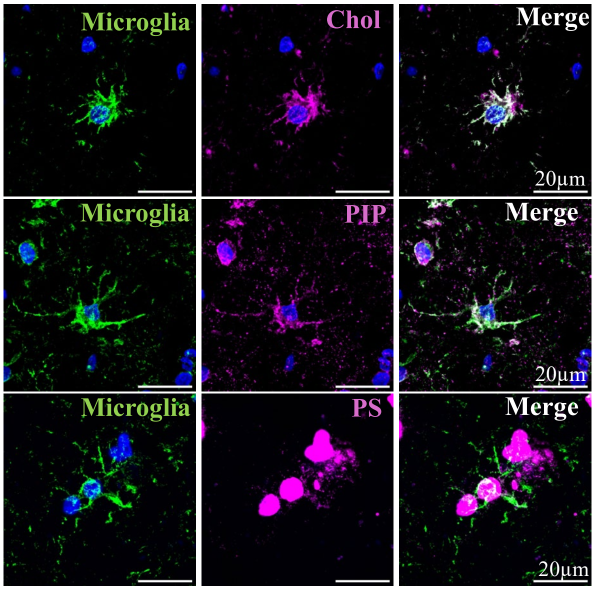


**Supplementary Figure 1: Anti-lipid antibodies reveal the spatial association of distinct lipid pools with microglia in fresh-frozen postmortem human AD sections.** Immunofluorescence profiling of distinct lipid targets within microglia. Representative images display co-staining of the microglia marker IBA1 (green) alongside specific lipids (magenta): cholesterol (top row), PIP (middle row), and PS (bottom row). Merged images (rightmost) highlight areas of colocalization (white) between microglia and each respective lipid. Scale bars, 20 µm.

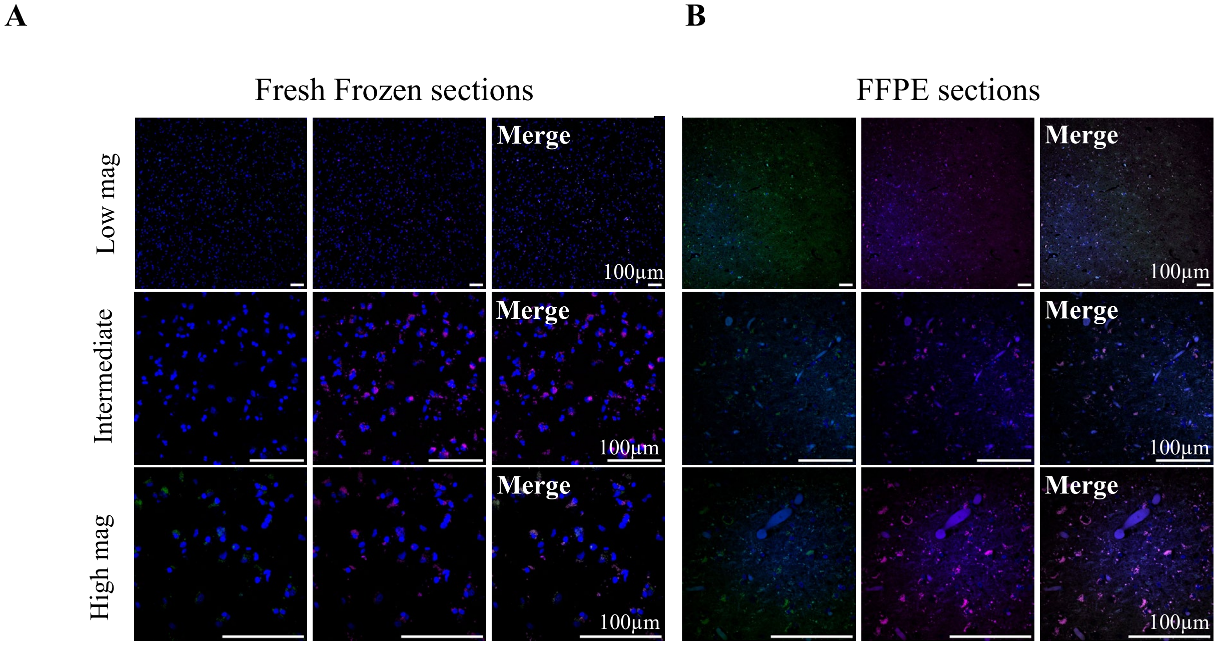
A
B
Fresh Frozen sections
FFPE sections
Low mag
Intermediate
High mag
Merge
100μm

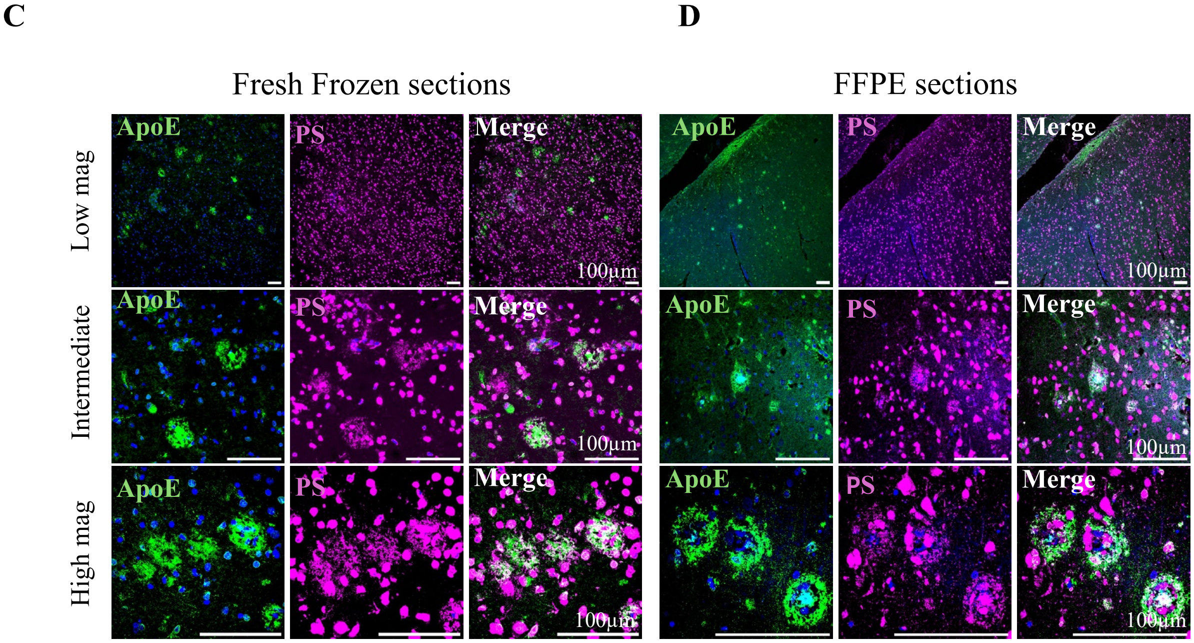
C
D
Fresh Frozen sections
FFPE sections
Low mag
Intermediate
High mag
ApoE
PS
Merge
100μm

10 **Supplementary Figure 2: Secondary antibody-only negative controls and preservation of phosphatidylserine in human postmortem AD brain sections.** Confocal microscopy assessing assay specificity and the spatial distribution of ApoE and phosphatidylserine (PS) in lipid-preserved (fresh-frozen) versus lipid-depleted (FFPE) postmortem human AD brain sections. Within each condition, images denote the green channel (left), the magenta channel (middle), and the merged channel (white, right). Rows represent increasing magnifications from top (low power) to bottom (high power). **(A)** Secondary antibody-only controls in fresh-frozen sections. Tissue sections were incubated without primary antibodies to assess non-specific background autofluorescence. **(B)** Secondary antibody-only controls in FFPE sections. **(C)** Co-staining for ApoE (green) and PS (magenta) in fresh-frozen sections. **(D)** Co-staining for ApoE (green) and PS (magenta) in FFPE sections. Consistent with findings for other anionic lipids, PS immunoreactivity remains highly preserved across both fresh-frozen and FFPE preparations. Scale bars, 100 µm.

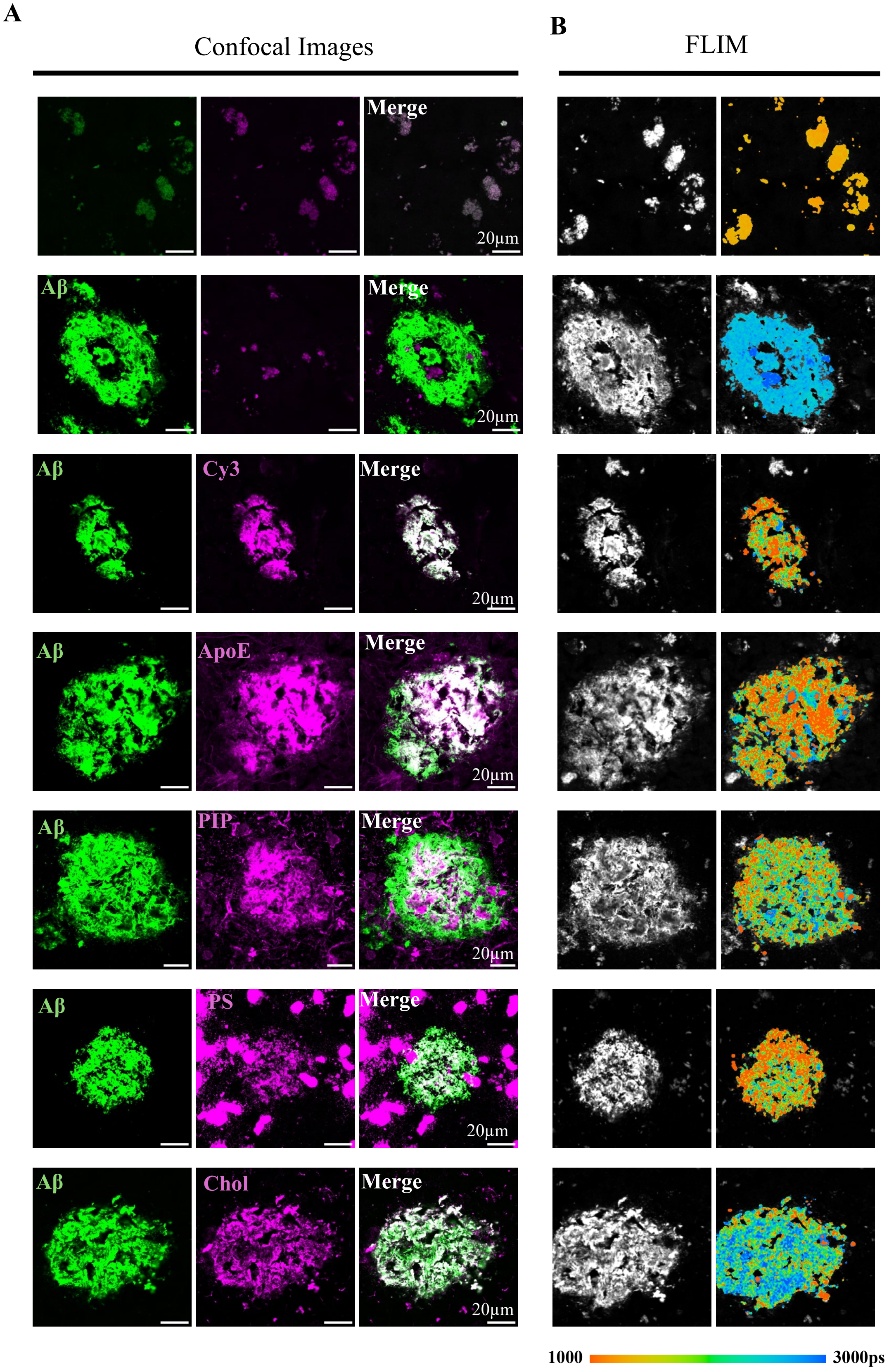
A
Confocal Images
B
FLIM
Merge
20μm
Aβ
Merge
20μm
Aβ
Cy3
Merge
20μm
Aβ
ApoE
Merge
20μm
Aβ
PIP
Merge
20μm
Aβ
PS
Merge
20μm
Aβ
Chol
Merge
20μm
1000
3000ps

12 **Supplementary Figure 3: FLIM-FRET analysis reveals significant quenching of Aβ donor lifetimes by anionic lipids, cholesterol, and ApoE in human postmortem AD brain sections.** Standard image layout for all experimental conditions. (**A**) The first three columns display conventional confocal imaging of the Aβ donor (green), the indicated Cy3 acceptor (magenta), and the merged channels (white). (**B**) Subsequent columns display the corresponding FLIM intensity maps and pseudo-colored FLIM lifetime images. Experimental conditions and established FRET pairs. *Top Row*: Unstained human AD tissue to measure intrinsic autofluorescence and establish the baseline reference lifetime. *Second Row*: Aβ donor-only condition establishing the unquenched baseline fluorescence lifetime. *Third Row*: Positive FRET control utilizing Aβ labeled with a secondary antibody dual-tagged with both A488 and Cy3 to demonstrate maximal lifetime quenching. *Rows 4–7*: Experimental FRET conditions demonstrating close nanometer-scale interactions between the Aβ donor and distinct Cy3-labeled acceptors: ApoE (fourth row), PIP (fifth row), PS (sixth row), and cholesterol (bottom row). Scale bars, 20 μm. Quantified fluorescence lifetimes (picoseconds) derived from these images (brain donor ID 2219) are summarized in Supplementary Table 2.

**Supplementary Table 2: Summary of Aβ-A488 donor lifetimes (ps) across various FRET-acceptor pairs.**

| | *APOEε4* homozygotes | | | *APOEε3* homozygotes | | |
|---|---|---|---|---|---|---|
| | **2219** | **1707** | **1967** | **2302** | **2594** | **2207** |
| **Cy3-labeled Acceptor** | | | | | | |
| D54D2 Aβ (donor only) | 2592 ± 22 | 2330 ± 25 | 2350 ± 58 | 2371 ± 33 | 2536 ± 32 | 2361 ± 26 |
| Positive Control | 1418 ± 186 | 868 ± 47 | 1141 ± 37 | 1174 ± 25 | 1718 ± 86 | 1033 ± 35 |
| ApoE | 1379 ± 157 | 1843 ± 93 | 1586 ± 107 | 1329 ± 160 | 1618 ± 80 | 1230 ± 71 |
| PIP | 1755 ± 147 | 1926 ± 108 | 1701 ± 82 | 1730 ± 294 | 1899 ± 90 | 2147 ± 63 |
| PS | 1398 ± 166 | 1387 ± 107 | 1894 ± 80 | 2000 ± 105 | 2008 ± 99 | 1904 ± 105 |
| Cholesterol | 2236 ± 138 | 1636 ± 74 | 1647 ± 124 | 1632 ± 95 | 1346 ± 103 | 1626 ± 121 |
| *6E10 Aβ (donor only) | 2570 ± 12 | 2374 ± 17 | 2418 ± 31 | 2508 ± 28 | 2649 ± 22 | 2498 ± 65 |

**Supplementary Table 2: Fluorescence lifetimes of Aβ-A488 donors.** Lifetimes (ps) are reported for donor-only controls, positive controls, and experimental groups paired with Cy3-labeled acceptors. Data are presented as mean lifetime values ± SD derived from independent experimental samples (N = 3). Statistical significance ($p < 0.0001$ for all samples) was determined by comparing donor lifetimes in FRET pairs against donor-only controls, with analysis performed on 10 Regions of Interest (ROIs) per sample. *The 6E10 Aβ-A488 antibody was used as the donor only pair for ApoE-Cy3 FRET to prevent cross-species contamination; all other interactions in this table were referenced to the D54D2 Aβ-A488 control.

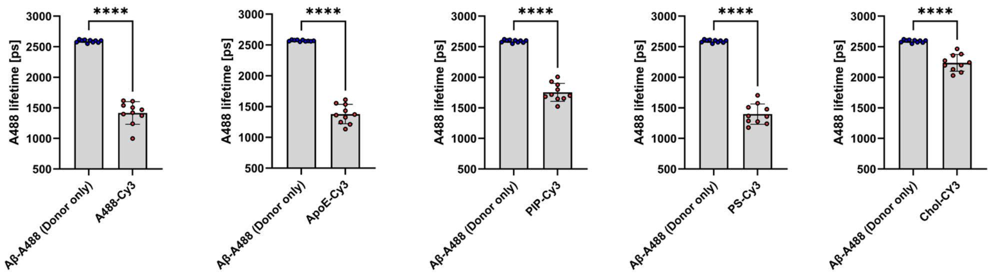


**Supplementary Figure 4: FLIM-FRET quantification of Aβ nanoscale proximity to lipid pools and ApoE in human AD brain sections.** Quantification of Aβ-A488 donor fluorescence lifetimes. Bar graphs compare the unquenched baseline lifetime of the Aβ-A488 donor against five specific Cy3-labeled acceptor conditions: (1) dual-labeled positive control, (2) ApoE, (3) PIP, (4) PS, and (5) cholesterol. Significant reductions in donor lifetime indicate nanoscale intermolecular proximity. For all panels, error bars represent standard deviation (s.d.). Statistical significance was determined using t-test with Welch's correction for uneven variances (****$p < 0.0001$).
